# Macrophage subtypes inhibit breast cancer proliferation in culture

**DOI:** 10.1101/2024.06.01.596963

**Authors:** Sophia R.S. Varady, Daniel Greiner, Minna Roh-Johnson

## Abstract

Macrophages are a highly plastic cell type that adopt distinct subtypes and functional states depending on environmental cues. These functional states can vary wildly, with distinct macrophages capable of displaying opposing functions. We sought to understand how macrophage subtypes that exist on two ends of a spectrum influence the function of other cells. We used a co-culture system with primary human macrophages to probe the effects of macrophage subtypes on breast cancer cell proliferation. Our studies revealed a surprising phenotype in which both macrophage subtypes inhibited cancer cell proliferation compared to cancer cells alone. Of particular interest, using two different proliferation assays with two different breast cancer cell lines, we showed that differentiating macrophages into a “pro-tumor” subtype inhibited breast cancer cell proliferation. These findings are inconsistent with the prevailing interpretation that “pro-tumor” macrophages promote cancer cell proliferation and suggest a re-evaluation of how these interpretations are made.

## Introduction

Macrophages are a component of the innate immune system, known for their role in killing microorganisms and removing dead cells. What is perhaps less known is that macrophages are a highly flexible cell type found in almost all tissues (for review, see (Mosser and Edwards, 2008)). Distinct macrophage differentiation states lead to their functional diversity, in which macrophages regulate a variety of processes including development (Eom and Parichy, 2017), cancer progression (Lin and Pollard, 2004; Nielsen and Schmid, 2017), and repair of blood vessels (Liu *et al*., 2016; Gurevich *et al*., 2018). Thus, macrophages are a unique cell type, in that an individual cell can transition between many different phenotypic states within its lifetime (Azizi *et al*., 2018; Locati *et al*., 2020; Cheng *et al*., 2021b).

Macrophages are often classified into two major subtypes representing two distinct ends of an activation spectrum: M1-activated or M2-activated (Ginhoux *et al*., 2016; Shapouri-Moghaddam *et al*., 2018; Yunna *et al*., 2020; Boutilier and Elsawa, 2021). M1-like macrophages are classically activated, pro-inflammatory macrophages, known for their role in eliminating pathogens and their anti-tumorigenicity (Sica and Mantovani, 2012; Martinez and Gordon, 2014; Pan *et al*., 2020). Conversely, M2-like macrophages are anti-inflammatory and function in tissue healing and tumor growth (Mantovani *et al*., 2002; Shapouri-Moghaddam *et al*., 2018; Boutilier and Elsawa, 2021). Both macrophage subtypes are typically classified by their surface marker expression, but these two classifications are simplistic and recent studies have revealed variation in the expression of these markers (Ginhoux *et al*., 2016; Muller *et al*., 2017; Azizi *et al*., 2018).

M2-like macrophages are representative of macrophages in the tumor (Hughes *et al*., 2015; Steenbrugge *et al*., 2018). Macrophages are abundant in many solid tumors and promote cancer progression (Lin and Pollard, 2004; Chen *et al*., 2005; Obeid *et al*., 2013; Mantovani *et al*., 2017; Nielsen and Schmid, 2017; Hinshaw and Shevde, 2019; Pan *et al*., 2020; Xiang *et al*., 2021). The existing paradigm is that M1-like macrophages inhibit cancer growth, and M2-like macrophages promote cancer growth (Zhang *et al*., 2014; Liu *et al*., 2021; Zhang and Sioud, 2023). However, there are a number of factors that complicate interpretations. These conclusions are based on comparisons of cancer cell proliferation in culture with M1-like *versus* M2-like macrophages (Sousa *et al*., 2015; Moraes *et al*., 2017; Zhang *et al*., 2023a), but do not reveal how macrophage subtypes regulate cancer cell proliferation compared to cancer cells alone (Sousa *et al*., 2015; Moraes *et al*., 2017; Zhang *et al*., 2023a). These comparisons are critical to interpret how macrophages ultimately regulate cancer cell proliferation. Furthermore, it is important to note that most of the reported studies use macrophage conditioned media approaches (Mu *et al*., 2018; Deng *et al*., 2021; Zhang *et al*., 2021; Chen *et al*., 2022; Cheng *et al*., 2023; Nie *et al*., 2023). While informative, these studies do not address how cell contact between macrophages and cancer cells regulate outcomes. Finally, many approaches use macrophage cell lines (THP-1, U937 and RAW 264.7) which are cancer cells themselves (Chanput *et al*., 2014; Taciak *et al*., 2018; Nascimento *et al*., 2022), and there is a dearth of studies using physiologically relevant macrophages.

Given that macrophages adopt differentiation states that represent two ends of a spectrum, we sought to use this simplified system to fundamentally understand how different macrophage subtypes regulate the behavior of another cell. By comparing macrophage/cancer cell co-cultures to cancer cells alone, and taking advantage of a direct co-culture system, as well as the commonly employed conditioned media system, we made a surprising observation in which the presence of either M1-like or M2-like macrophages *inhibit* cancer cell proliferation compared to cancer cells alone. These results are in contrast to interpretations made from previous studies, and provide a previously undescribed function for M2-like macrophages in breast cancer proliferation.

## Results and Discussion

### M2-like macrophages do not promote cancer cell proliferative capacity

Previous literature has shown that macrophages differentiated into an M2-like subtype promote cancer cell proliferation when compared to macrophages differentiated into an M1-like subtype (Liu *et al*., 2020; Zhao *et al*., 2021; Zhang *et al*., 2023a). These findings are consistent with data generated *in vivo*, in which tumor-associated macrophages are tumor-promoting (Dandekar *et al*., 2011; Cendrowicz *et al*., 2021; Munir *et al*., 2021), although we recognize that the complexity of tumor-associated macrophages is not recapitulated with *in vitro* macrophage polarization studies (Jia *et al*., 2016; Ramesh *et al*., 2020; Ma *et al*., 2022). Nevertheless, these findings have put forth a model by which M2-like macrophages promote cancer cell proliferation. To understand how macrophage subtypes regulate cancer cell proliferation compared to cancer cells alone, we used an *in vitro* system using primary human macrophages (Figure 1A). We isolated primary monocytes from healthy blood donors, and differentiated them into either an M1-like or M2-like macrophage state using previously published approaches (Jia *et al*., 2016; Zheng *et al*., 2018; Little *et al*., 2019; Mohd Yasin *et al*., 2022; Kidwell *et al*., 2023) for subsequent experiments.

**Figure 1.**
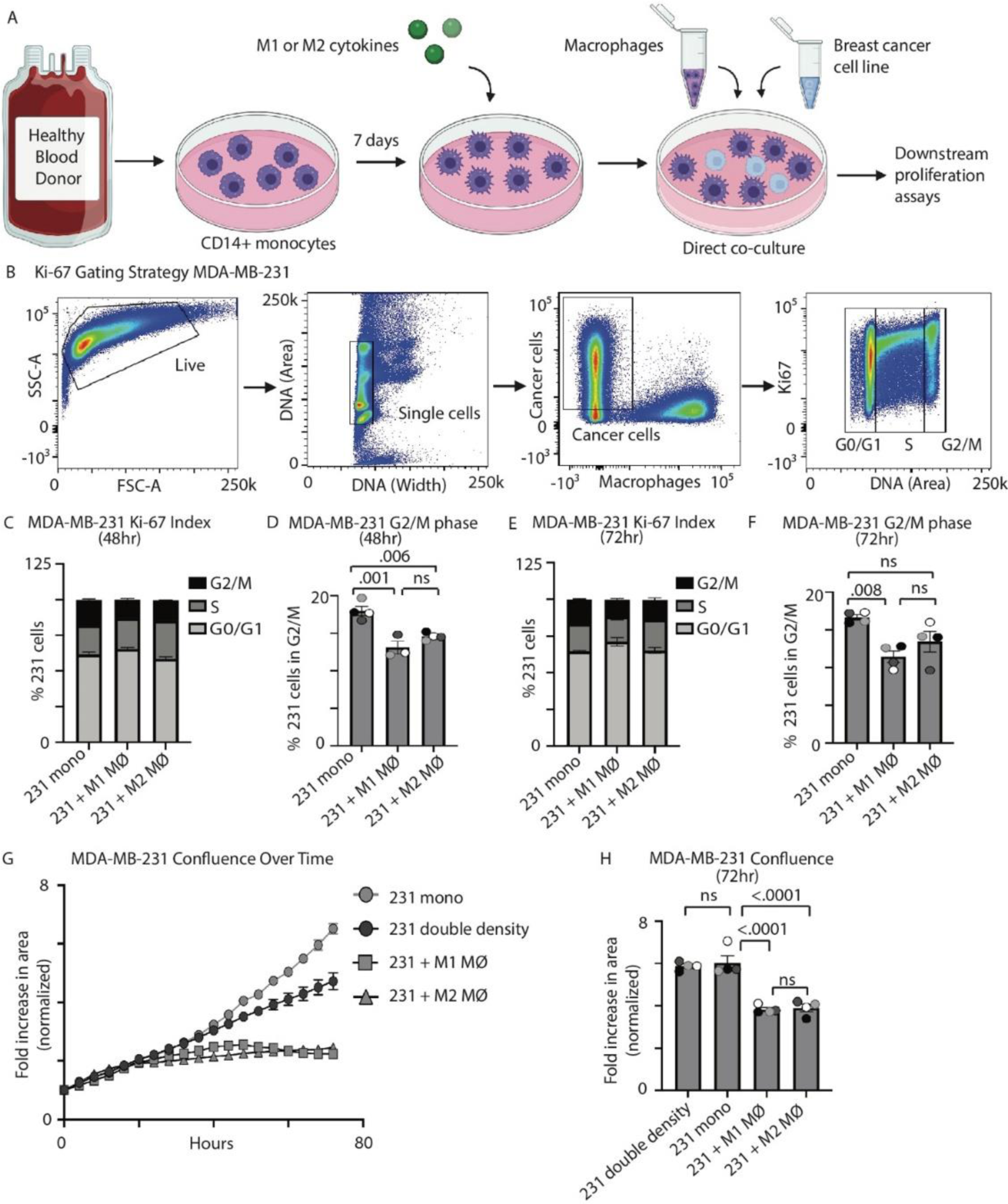
MDA-MB-231 breast cancer cells do not exhibit increased proliferation when in culture with M2-like macrophages. (A) Schematic outlining co-culture approaches. (B) Example gating of Ki67 cell cycle index flow cytometry assays. (C-F) Quantification of MDA-MB-231 cells in each stage of the cell cycle at 48 (C) and 72 (E) hours, following gating strategies outlined in (B) when MDA-MB-231 cells are plated alone (“231 mono”), when MDA-MB-231 cells are plated with M1-like macrophages (“231 + M1 MΦ”) and when MDA-MB-231 cells are plated with M2-like macrophages (“231 + M2 MΦ”) (n=4 independent donors). (D, F) Comparison of only the G2/M phase of the cell cycle from data in (C) at 48 hours and data in (E) at 72 hours (n=4; Tukey’s 1-way ANOVA). (G) Representative example of MDA-MB-231 cell confluence over 72 hours when MDA-MB-231 cells are plated under conditions described in (C), plus MDA-MB-231 cells plated alone at double density (“231 double density”). (H) Quantification of MDA-MB-231 cell confluence at 72 hours in (G), reported as the fold increase in cell area over time (n=4 independent donors; Tukey’s 1-way ANOVA).

M1-like or M2-like macrophages were plated in direct co-culture with a breast cancer cell line, MDA-MB-231 (231 cells), stably expressing a histone-localized mCherry (H2B-mCherry) (Figure 1A). 231 cells are a breast cancer line that lacks estrogen, progesterone, and HER2 receptors, and is thus referred to as “triple negative breast cancer”, which is a type of breast cancer with few treatment options and poor prognosis. We first used a flow cytometry assay to evaluate 231 proliferation changes by quantifying the presence of a proliferation marker Ki67 (Endl and Gerdes, 2000; Kim and Sederstrom, 2015), as well as DNA content. With this approach, we quantified the percentage of 231 cells within the G2 and Mitotic (M) phases of the cell cycle (Figure 1B). When comparing 231 proliferative capacity in the presence of M1-like macrophages versus cancer cells alone, 231 cells exhibited a significantly lower percentage of cells in the G2/M phase of the cell cycle, which is consistent with previous reports showing “anti-tumor” activity of M1-like macrophages (Figure 1, C-F) (Jia *et al*., 2016; Ramesh *et al*., 2020; Moradi-Chaleshtori *et al*., 2021). However, quite surprisingly, when comparing the percentage of cancer cells within the G2/M phase of the cell cycle when in co-culture with M2-like macrophages versus cancer cells alone, we also observed a significant decrease in cancer cell proliferation when cancer cells were cultured with M2-like macrophages at 48 hours (Figure 1, C-D), which continued to trend at 72 hours (Figure 1, E-F). These results were surprising given that the prevailing model is that M2-like macrophages promote cancer cell proliferation (Liu *et al*., 2020; Chen *et al*., 2021; Zhang *et al*., 2023a).

The increase in the population of cancer cells within the G2/M phase of the cell cycle could reflect increased proliferation or a G2/M cell cycle arrest. Thus, we took advantage of an assay that quantifies cell confluence over time (Figure 1, G-H). Consistent with the cell cycle results, we observed a significant decrease in cancer cell proliferation when cancer cells are co-cultured with either M1-like or M2-like macrophages (Figure 1, G-H; Figure S1A; Supplemental Video S1) compared to cancer cells alone. To rule out the effects of cell confluence, we plated cancer cells at the same number as that used in co-cultures (231 mono) as well as double the number (231 double density) and found no difference in cancer cell proliferation between these two conditions (Figure 1, G-H). Furthermore, we asked whether the culture system may be masking a proliferation phenotype if the cancer cells are already dividing at maximum capability. Thus, we performed the same experiments in lower serum and first found that the basal proliferation rate of 231 cells was reduced when cells were cultured in 0.5% and 2% serum compared to full serum (10%) (Figure S1B). Using these low serum conditions, consistent with our previous results, M2-like macrophages in both 0.5% and 2% serum still inhibited breast cancer cell proliferation compared to cancer cells alone at 72 hours (Figure S1, C-D). These results suggest that M2-like macrophages do not promote breast cancer cell proliferation.

### M2-like macrophages retain their M2 status over time

Due to the surprising results that M2-like macrophages do not promote cancer cell proliferation compared to cancer cells alone, we sought to confirm the differentiation status of the macrophages. We hypothesized that the lack of cancer cell proliferation in the presence of M2-like macrophages could be due to macrophages losing their M2-like state over time (see methods). We performed flow cytometry analysis of CD206 expression (M2-like marker) (Figure 2, A-C; Figure S2, A-B) as has been done previously (Shrivastava *et al*., 2019; Li *et al*., 2024; Tang *et al*., 2024; Zhang *et al*., 2024). M2-like macrophages in co-culture with 231 cells maintain expression of CD206 from 24-72 hours at a similar level to control (M2-like macrophages alone) (Figure 2, A-C) We also quantified M1-like macrophage status over time using CD80 (M1-like marker). We found that M1-like macrophages exhibited higher levels of CD80/CD206 expression at 24, 48 and 72 hours compared to M2-like macrophages as has been done previously (Ramesh *et al*., 2020) (Figure S2, C-F). These results suggest that both M1-like and M2-like macrophages retain their differentiation status during the course of the experiments.

**Figure 2.**
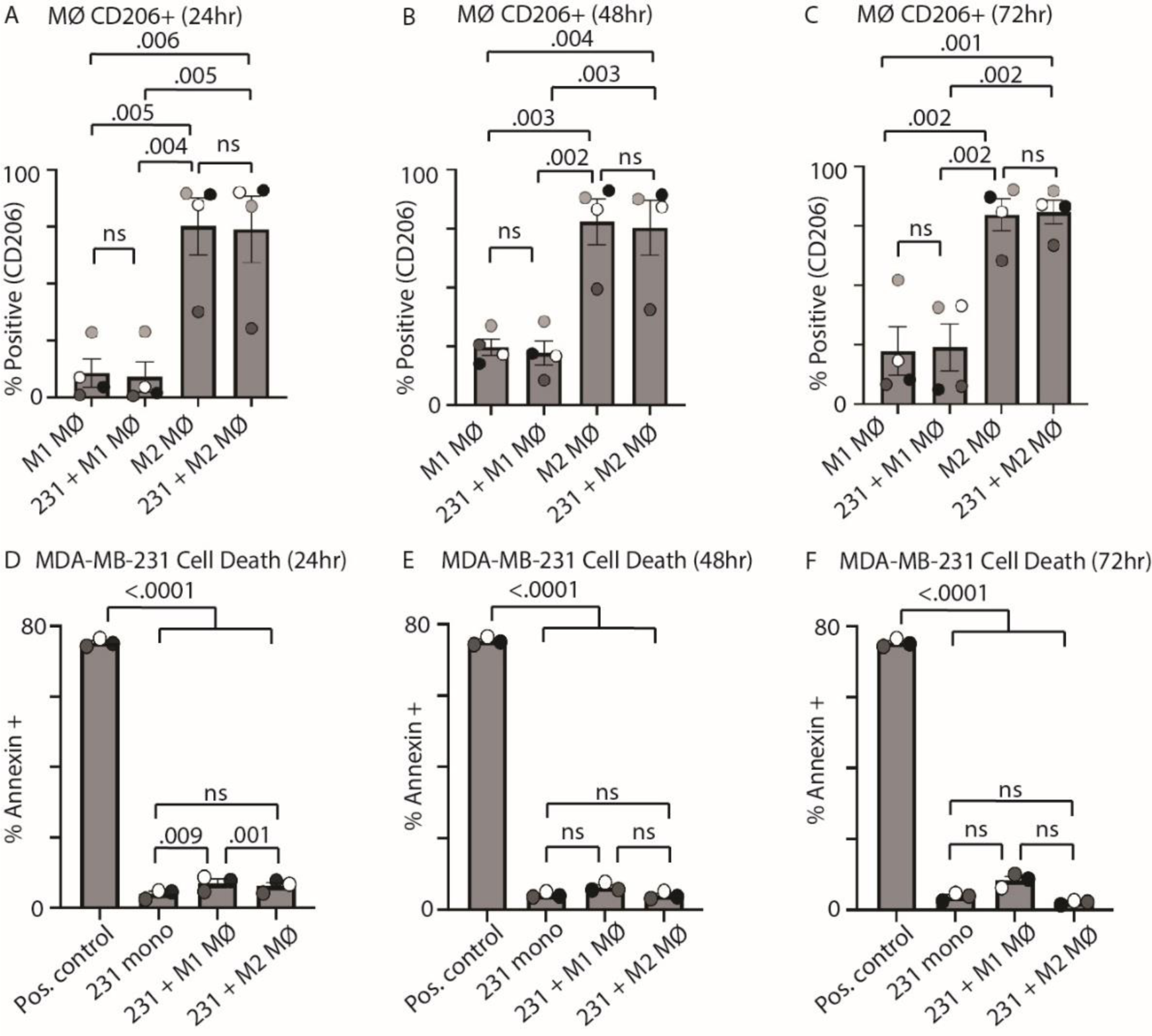
M2-like macrophages maintain differentiation state over the experimental time course and do not induce MDA-MB-231 cell death. (A-C) Quantification of CD206+ macrophages across culture conditions. Quantification of the percent of CD206+ macrophages measured by surface CD206 expression at 24 hours (A), 48 hours (B), and 72 hours (C) of culture when M1-like macrophages are plated alone (“M1 MΦ”), when M1-like macrophages are plated with MDA-MB-231 cells (“231 + M1 MΦ”), when M2-like macrophages are plated alone (“M2 MΦ”), and when M2-like macrophages are plated with MDA-MB-231 cells (“231 + M2 MΦ”) (n=4 independent donors; Tukey’s 1-way ANOVA). (D-F) Quantification of Annexin V positive MDA-MB-231 cells at 24 hours (D), 48 hours (E), and 72 hours (F) when MDA-MB-231 cells are plated as described in (A-C). The positive control is heat-killed cells (see methods) (n=3 independent donors; Tukey’s 1-way ANOVA). Analysis was performed on the same day; thus, the same positive control was used for all time points.

### Lack of cancer cell proliferation with M2-like macrophages is not due to increased cancer cell death or spreading

We next determined whether the reduced breast cancer cell number observed with M2-like macrophages was due to reduced cancer cell proliferation or increased cancer cell death. We quantified cancer cell death via Annexin V staining via flow cytometry (Figure 2, D-F; Figure S3, A-C). Annexin V detects phosphatidylserine, which is indicative of early stages of apoptosis (van Engeland *et al*., 1998). We found that at 24 hours, M1-like macrophages increased 231 cell death compared to 231 cells alone and M2-like macrophage co-culture (Figure 2D). At 48 and 72 hours, cancer cells in culture with M1-like macrophages exhibited subtle, but non-significant, increases in cancer cell death compared to cancer cells alone and cancer cells co-cultured with M2-like macrophages (Figure 2, E-F). This increase in cancer cell death in the presence of M1-like macrophages is consistent with previous reports that M1-like macrophages can be cytotoxic(Italiani and Boraschi, 2014; Shapouri-Moghaddam *et al*., 2018; Najafi *et al*., 2019). However, the level of cancer cell death observed in this condition (8.3% at 72 hours) does not account for the decreased cancer cell proliferation observed in our cancer cell + M1-like macrophage co-cultures (37% decrease) (Figure 1H). Furthermore, 231 cells in co-culture with M2-like macrophages did not show differences in cell death compared to 231 cells alone (Figure 2, D-F), suggesting the observed decrease in cancer cell proliferation in the presence of M2-like macrophages is not due to cell death.

We measured proliferation in the time courses by quantifying the amount of surface area on the plate taken up by cancer cells over time (Figure 1, G-H). Thus, it is possible that increases in overall surface area are due to increased cancer cell spreading and not increases in cell number. To test between these hypotheses, we measured individual cancer cell area at 12 hours (time of cell attachment) and 72 hours (final time point) (Figure S3, D-E). At 12 hours, we did not observe any differences in cancer cell area across conditions (Figure S3D). At 72 hours, cancer cells exhibited a smaller cell area when in co-culture with M1-like macrophages (Figure S3E); however, importantly, we did not observe a difference in cell area of cancer cells in culture with M2-like macrophages versus cancer cells alone. Taken together with our cell death analysis, these results support the model that cancer cell proliferation is inhibited in the presence of M2-like macrophages.

### M2-like macrophages do not promote cancer cell proliferation in MDA-MB-468 cells

Our findings that M2-like macrophages do not promote breast cancer cell proliferation were unexpected. Thus, we tested this hypothesis in another triple negative breast cancer cell line, MDA-MB-468 (468 cells), under the same culture conditions. Using flow cytometry to quantify Ki67 and DNA content, M2-like macrophages did not promote 468 cell proliferation at 48 and 72 hours (Figure 3, A-D) compared to cancer cells alone. We then quantified 468 cell proliferation over time and observed a decrease when in the presence of M2-like macrophages compared to 468 cells alone (Figure 3, E-F), consistent with results we observed with 231 cells (Figure 1H). Thus, these data further support our finding that M2-like macrophages do not promote cancer cell proliferation.

**Figure 3.**
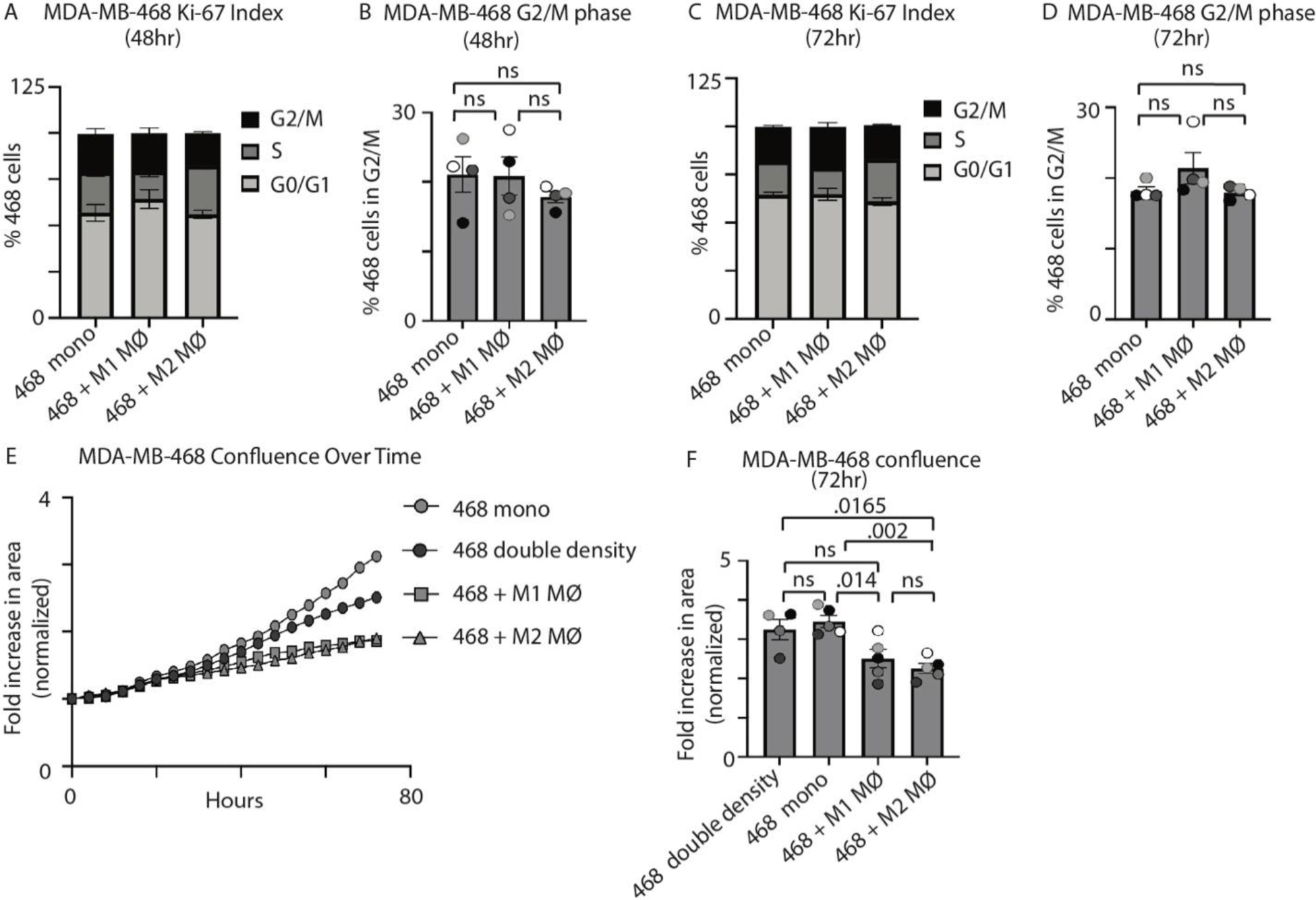
MDA-MB-468 cells do not show increased proliferation when in culture with M2-like macrophages. (A-D) Quantification of MDA-MB-468 cells in each stage of the cell cycle at 48 (A) and 72 (C) hours when MDA-MB-468 cells are plated alone (“468 mono”), when MDA-MB-468 cells are plated with M1-like macrophages (“468 + M1 MΦ”) and when MDA-MB-468 cells are plated with M2-like macrophages (“468 + M2 MΦ”) (n=4 independent donors). (B, D) Comparison of only the G2/M phase of the cell cycle from data in (A) at 48 hours and data in (C) at 72 hours (n=4; Tukey’s 1-way ANOVA). (E) Representative example of MDA-MB-468 cell confluence over 72 hours when MDA-MB-468 cells are plated as in (A), plus MDA-MB-468 cells are plated alone at double density (“468 double density). (F) Quantification of MDA-MB-468 cell confluence at 72 hours in (E), reported as the fold increase in cell area over time (n=4 independent donors; Tukey’s 1-way ANOVA).

### Conditioned media from M2-like macrophages do not promote cancer cell proliferation

The discrepancy in our results compared to the field caused us to compare our approaches with conditioned media approaches used previously (Mor *et al*., 1998; Sousa *et al*., 2015; Jedrzejewski *et al*., 2020; Li *et al*., 2020). We used conditioned media from either M1-like or M2-like macrophages alone (M1 mono CM or M2 mono CM), as well as conditioned media from macrophage/cancer cell co-cultures (ie. M1 + 231 CM or M2 + 231 CM), and asked how each of these conditioned media affects the proliferation of breast cancer cells. We found that conditioned media from M2-like macrophages alone (Figure 4, A-B) or 231 + M2 MΦ co-culture (Figure 4C) caused subtle increases in 231 cell proliferation versus M1 conditioned media, although not statistically significant. With 468 cells, we found that 468 cells in M2 + 468 conditioned media exhibited significantly increased proliferation compared to M1 + 468 conditioned media at 72 hours (Figure 4, D-F). Together these results suggest that when comparing the effects of M2-like macrophages on cancer cell proliferation to M1-like macrophages, we observe results that are consistent with those in the field (Zhang *et al*., 2021; Zhao *et al*., 2021; Cheng *et al*., 2023).

**Figure 4.**
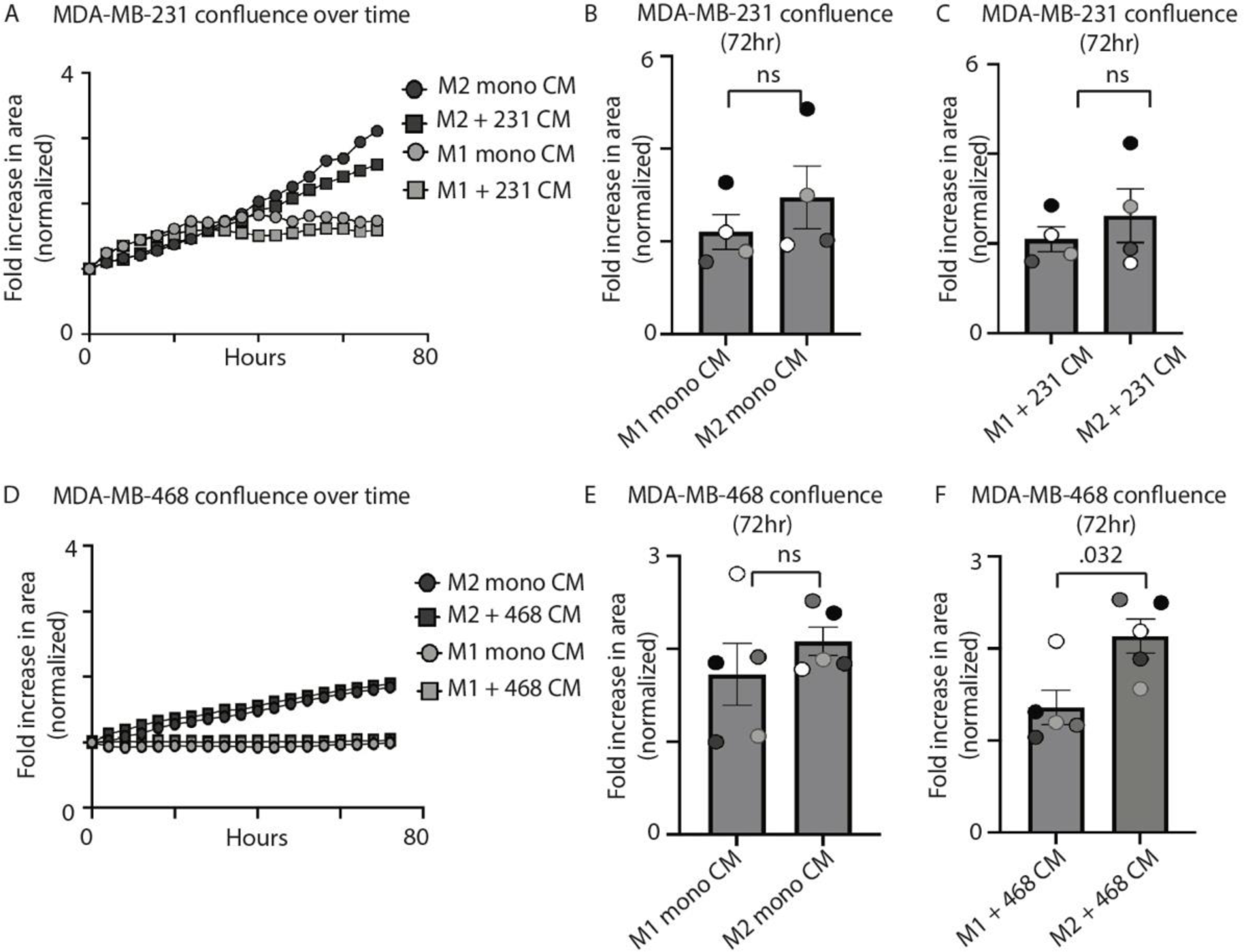
Conditioned media from M2-like macrophages increases MDA-MB-468 proliferation compared to conditioned media from M1-like macrophages. (A) Representative example of proliferation of MDA-MB-231 cell confluence over 72 hours when naïve MDA-MB-231 cells are treated with M2-like macrophage conditioned media (“M2 mono CM”), when naïve MDA-MB-231 cells are treated with MDA-MB-231/M2-like macrophage co-culture conditioned media (“M2 + 231 CM”), when naïve MDA-MB-231 cells are treated with M1-like macrophage conditioned media (M1 mono CM), and when naïve MDA-MB-231 cells are treated with MDA-MB-231/M1-like macrophage co-culture conditioned media (“M1 + 231 CM”). (B) Quantification of MDA-MB-231 cell confluence at 48 hours when treated with M1-like or M2-like macrophage monoculture CM in (A) (n=4 independent donors; Mann-Whitney test). (C) Quantification of MDA-MB-231 cell confluence at 72 hours when treated with MDA-MB-231/M1-like or MDA-MB-231/M2-like macrophage co-culture conditioned media in (A) (n=5 independent donors; Mann-Whitney test). (D) Representative example of proliferation of MDA-MB-468 cell numbers over 72 hours when naïve MDA-MB-468 cells are treated as described in (A). (E) Quantification of MDA-MB-468 cell confluence at 48 hours when treated with M1-like or M2-like macrophage monoculture CM in (D) (n=5 independent donors; Mann-Whitney test). (F) Quantification of MDA-MB-468 cell confluence at 72 hours when treated with MDA-MB-468/M1-like or MDA-MB-468/M2-like macrophage co-culture conditioned media in (D) (n=5 independent donors; Mann-Whitney test).

We next asked how macrophage conditioned media regulates cancer cell proliferation versus cancer cells alone. Additionally, we asked how these conditions compare to direct contact co-cultures. We first found that the presence of M1-like macrophages (either in direct co-cultures or conditioned media experiments) inhibited 231 cell proliferation compared to 231 cells alone (Figure 5, A-B), again consistent with previous reports of M1-like macrophage-induced cytotoxicity (Engstrom *et al*., 2014; Moradi-Chaleshtori *et al*., 2021). Interestingly, M1 macrophage conditioned media inhibited cancer cell proliferation better than direct 231 + M1 MΦ co-cultures (Figure 5, A-B). However, we found that conditioned media from M2-like macrophages did not increase cancer cell proliferation compared to cancer cells alone (Figure 5, C-D), as previous literature has suggested (Deng *et al*., 2021; Moradi-Chaleshtori *et al*., 2021; Cheng *et al*., 2023; Nie *et al*., 2023; Yu *et al*., 2023; Zhang *et al*., 2023a). When we performed the same experiments with 468 cells, we observed results that were consistent with the 231 experiments (Figure 5, E-H). We again tested the effects of macrophage conditioned media on cancer cell proliferation under low serum conditions. In low serum conditions (0.5% and 2%), conditioned media from M2-like macrophages, either from M2-like macrophages alone or 231 + M2 MΦ co-cultures, did not promote 231 cell proliferation compared to 231 cells alone (Figure S4, A-B).

**Figure 5.**
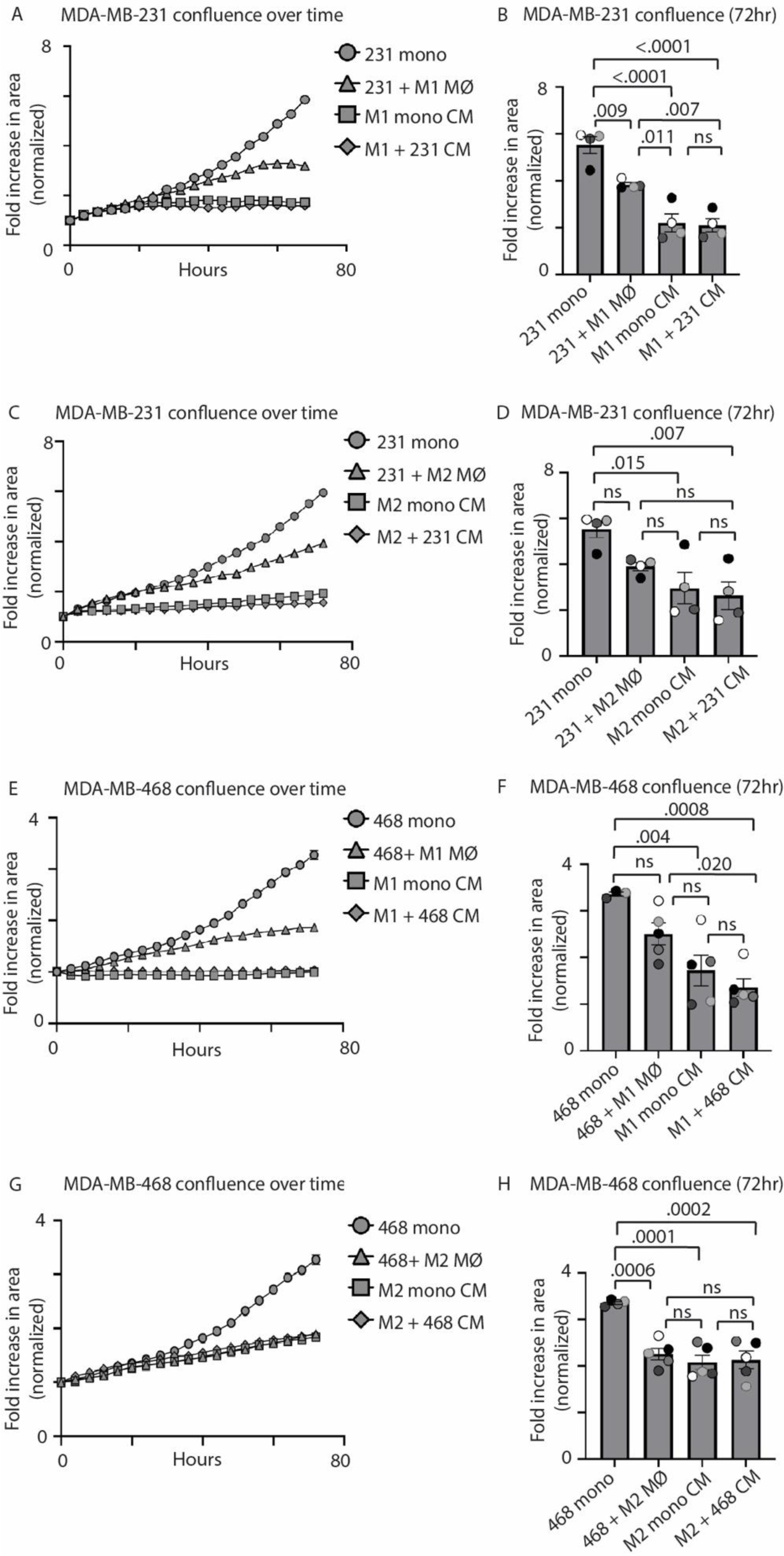
Conditioned media from M2-like macrophages inhibit breast cancer cell proliferation compared to cancer cells alone. (A) Representative example of proliferation of MDA-MB-231 cell numbers over 72 hours when naïve MDA-MB-231 cells are treated with MDA-MB-231 monoculture conditioned media (“231 mono”), when MDA-MB-231 cells are in direct co-culture with M1-like macrophages (“231 + M1 MΦ”), when naïve MDA-MB-231 cells are treated with MDA-MB-231/M1-like macrophage co-culture conditioned media (“M1 mono CM”), and when naïve MDA-MB-231 cells are treated with MDA-MB-231/M1-like macrophage co-culture conditioned media (“M1 + 231 CM”). (B) Quantification of MDA-MB-231 cell confluence at 72 hours when plated in conditions listed in (A) (n=4 independent donors; Tukey’s 1-way ANOVA). (C) Representative example of proliferation of MDA-MB-231 cell numbers over 72 hours when naïve MDA-MB-231 cells are treated with conditions described in (A), but with M2 macrophages. (D) Quantification of MDA-MB-231 cell confluence at 72 hours when plated in conditions listed in (C) (n=4 independent donors; Tukey’s 1-way ANOVA). (E) Representative example of proliferation of MDA-MB-468 cell numbers over 72 hours when naïve MDA-MB-468 cells are treated with conditions described in (A). (F) Quantification of MDA-MB-468 cell confluence at 72 hours when plated in conditions listed in (E) (n=5 independent donors; Tukey’s 1-way ANOVA). (G) Representative example of proliferation of MDA-MB-468 cell numbers over 72 hours when naïve MDA-MB-468 cells are treated with conditions described in (B). (H) Quantification of MDA-MB-468 cell confluence at 72 hours when plated in conditions listed in (G) (n=5 independent donors; Tukey’s 1-way ANOVA).

While many studies have investigated the role of macrophage subtypes on cancer progression, comparisons are typically made *between* macrophage subtypes, leading to conclusions that M2-like macrophages promote cancer cell proliferation (Liu *et al*., 2020; Cheng *et al*., 2023; Hao *et al*., 2023; Yu *et al*., 2023). However, using direct co-culture systems, conditioned media approaches, and two proliferation assays, our study shows an unexpected phenotype in which M2-like macrophages do not promote breast cancer cell proliferation compared to breast cancer cells alone, and may, in fact, decrease breast cancer cell proliferation. These results raise several interesting points about how macrophages regulate cancer cell behavior, and how this process relates to cancer progression.

The small number of studies that have compared cancer cell proliferation when in culture with macrophages versus cancer cells alone present mixed results, with studies showing that M2-like macrophages promote breast cancer cell proliferation (Carroll *et al*., 2016; Hao *et al*., 2023), and others showing that M2-like macrophages inhibit breast cancer cell proliferation (Lindsten *et al*., 2017; Hirano *et al*., 2023). Our work is consistent with the results from the latter studies, and suggest that the different results obtained from these experiments could be due to the type of macrophage (primary versus cell line), how the macrophages were differentiated, and the cancer cell type. Additional work is required to fully understand this process.

There is a wealth of data showing that the presence of tumor-associated macrophages promote tumor growth and metastasis (Cheng *et al*., 2021a; Kumari and Choi, 2022; Zhang *et al*., 2023b; Zhao *et al*., 2024). How do we reconcile these results with our results showing that the presence of macrophages decreases breast cancer cell proliferation compared to cancer cells alone? In our study, we do not model tumor-associated macrophages since differentiating macrophages in culture will not fully recapitulate the macrophage differentiation status in a tumor. However, we used our system to fundamentally understand how “pro-tumor” M2-like macrophages versus “anti-tumor” M1-like macrophages specifically regulate cancer cell proliferation. Given that macrophage infiltration correlates with tumor grade, these results suggest that early in tumorigenesis (Mahmoud *et al*., 2012; Yuan *et al*., 2019; Jamiyan *et al*., 2020), cancer cells begin to proliferate in the absence of stromal cells. Indeed, macrophage infiltration is still an early step in breast cancer tumorigenesis (Zhukova *et al*., 2020; Zhu *et al*., 2021), but we would argue that whether tumor growth is a direct effect of pro-tumor macrophages or a consequence of the function of pro-tumor macrophages on other stromal cells is unclear. Tumor-associated macrophages have been shown to promote tumor angiogenesis which brings nutrients into the tumor, fueling tumor growth (Riabov *et al*., 2014; Fu *et al*., 2020). Tumor-associated macrophages also promote local cancer cell invasion and entry into the vasculature compared to cancer cells alone (Roh-Johnson *et al*., 2014; Harney *et al*., 2015). Thus, it is possible that the growth observed in tumors with either exogenous M2-like macrophages added (Cho *et al*., 2012; Steenbrugge *et al*., 2018) or differentiating endogenous macrophages into an M2-like status (Tao *et al*., 2020; Stachowicz-Suhs *et al*., 2024) is due to effects on the tumor environment, rather than directly on cancer cells. It is also possible that the direct role of tumor-associated macrophages on breast cancer cells is reduced cancer cell proliferation to support local cancer invasion, a key step in metastasis. Thus, future work is required to tease apart the direct functions of macrophages on cancer cell behavior.

## Acknowledgements

We would like to acknowledge Keren Hilgendorf, Joseph Casalini, and members of the Roh-Johnson lab for constructive feedback; and James Marvin, Meghan Curtin, and Elena Kurudza for technical support and advice. This work was supported by National Cancer Institute R37CA247994 to MRJ and 5P30CA042014-24 to the Flow Cytometry Core.

## Methods

### Cell culture of cell lines

Human cell lines MDA-MB-231 (HTB-26) and MDA-MB-468 (HTB-132) were purchased from the American Type Culture Collection (ATCC) and cultured at 37°C with 5% CO_2_ according to their recommendations. Base culture media used was DMEM (high-glucose; 11965118, ThermoFisher), and 10% heat-inactivated fetal bovine serum (FBS; F4135, ThermoFisher). All cell lines were kept in culture for no more than 20 passages total. The MDA-MB-468 cell line was supplemented with 1% sodium pyruvate. Mycoplasma testing was performed every 6 months using the Mycoplasma Detection Kit (30–1012 K, ATCC).

### Isolation and culture of PBMCs

Leukocyte cone filters were obtained from Associated Regional and Clinical Pathologists (ARUP) laboratories from unidentified healthy human blood donors. CD14+ monocytes were isolated as described(Greiner *et al*., 2022) with the exception of selection being adherence-based to isolate monocytes from buffy coats. Freshly harvested primary monocytes were plated at a density of 20-25 M per 10 cm plate in macrophage culture media containing: RPMI (11875119, ThermoFisher), 10% FBS (26140079, Thermo Fisher), 0.5% penicillin/streptomycin (P/S; P4333, Thermo Fisher), 10 mM HEPES (15630080, ThermoFisher), 0.1% 2-Mercaptoethanol (21985023, Thermo Fisher), and recombinant human GM-CSF at 20 ng/mL (300–03, Peprotech). The following day (day 1), additional macrophage media was added, and on day 4, macrophage media was replaced. On day 7, macrophages were differentiated into either M1 or M2 activation states, or remained undifferentiated as ‘M0’ like macrophages acting as an activation control.

### Macrophage activation by cytokines

To differentiate macrophages into an M1-like state, IFN-γ (3000–02, Peprotech, 50 ng/mL) and lipopolysaccharide (LPS; 00-4976-03, eBioscience, 10 ng/mL) were added to macrophage media for 72 hours prior to experiments. For M2-like activation, IL-4 (200-04, Peprotech, 20 ng/mL) and IL-13 (200–13, Peprotech, 20 ng/mL) were added to macrophage media for 72 hours prior to experiments. Once plated for experiments, both M1-like and M2-like macrophage media did not contain cytokines (macrophage culture media + GM-CSF only) so as not to effect cancer cells with activation cytokines.

### Cloning and cancer cell transduction

The pLenti6-H2B-mCherry plasmid was purchased from Addgene (89766). MDA-MB-231 and MDA-MB-468 cell lines were transduced with lentiviral plasmid vectors to create stable lines. First, HEK 293 FT cells were plated on poly-L-lysine-coated 15 cm dishes. HEK 293 FT cells were transfected with PEI-max (24765, Polysciences) and plasmids for pCMV-VSV-G, psPax2, and the H2B-mCherry transgene cassette. Cells were washed the following day and grown for an additional 36 hours in fresh media. The supernatants were harvested, passed through 0.45 μm filters, and used fresh as previously described (Johnson *et al*., 2020). Lentiviral supernatants were used to transduce cell lines by plating 50,000 cells into one well of a 6-well plate with the lentiviral supernatant, which was diluted 1:5 in DMEM media with a final concentration of 10 μg/mL polybrene (TR-1003-G, Sigma). After 48 hours, the cells were expanded and flow sorted using the BD FACS Aria cytometer to select for fluorescent expression.

### Ki67 cell cycle assay

MDA-MB-231 and MDA-MB-468 cell lines, respectively, were cultured directly with distinct macrophage subtypes (M1-like or M2-like, as described in ‘Isolation and culture of PBMCs’). Co-cultures for either cancer cell type were always cultured in macrophage media + GM-CSF (20 ng/mL) without additional polarization cytokines so as not to perturb cancer cells. Co-culture samples were enzymatically detached using Trypl-E Express Enzyme (1x) (12605028, ThermoFisher). Samples were incubated with staining buffer containing anti-human BV711-CD11b at 1:20 for 30 min on ice before fixation with the eBioscience Foxp3/Transcription Factor Staining Buffer Set (00-5523-00, ThermoFisher), according to manufacturer’s instructions. Cells were then stained with an APC conjugated Ki-67 antibody (APC-Ki67; 17-5699-42, ThermoFisher) at 1:20 for 30 min followed by a 3 µM DAPI (D9542, Sigma) solution for 10 min. Cells were resuspended in cold DPBS for analysis on the Fortessa Flow Cytometer. Single color controls were included for compensation in flow analysis. Single cells were selected by DNA width and DNA area, as has been done previously for Ki67 assays (Di Rosa *et al*., 2021; Kidwell *et al*., 2023), due to its increased specificity compared to standard single cell selection methods for flow cytometry. Cancer cells were distinguished from macrophages based on CD11b-negative controls and no color cancer cell controls. Gating strategies are outlined in Figure 1B.

### Incucyte cell proliferation assay and analysis

The Sartorius Incucyte S3 was used to measure MDA-MB-231 and MDA-MB-468 cell proliferation in various conditions including direct co-culture with M1-like or M2-like activated macrophages, as well as conditioned media conditions. Cell conditions were plated in replicates of 6 per experimental run and imaged at 20x magnification every 4 hours over the course of 72 hours. Proliferation was determined by cell area at each imaging time point. Cell area was defined by phase images and parameters were manually adjusted to correctly define individual cells. Total cell area per well was averaged across technical replicate wells of each condition. Confluence was reported as fold increase in phase area normalized to the first time point. The final time point (72 hours) fold increase in area was compared across conditions to measure statistical differences in changes in confluence. In addition, results were normalized to the first time point to account for slight variation in initial plating densities. Since macrophages were in the experimental system, but are post-mitotic, we included “double density” cancer cell monoculture controls to account for additional cell numbers in co-culture conditions, and confirm that initial confluence did not affect our results. For conditioned media experiments, naïve cancer cells were plated at a density of 7,500 cells/well in monoculture conditions (this includes all CM experiments) in a 96-well plate.

Cell spreading was assessed by measuring cell area of 231 cell monoculture cells, 231 cells in 231/M1-like macrophage co-cultures, and 231 cells in 231/M2-like macrophage co-cultures at 12 hours (when the cells first adhered to the plate) and 72 hours. Measurements were taken in ImageJ by manual cell selection and area measurement.

### Conditioned media experiments

Conditioned media conditions included M1 macrophage monoculture conditioned media, M1 co-culture with 231 or 468 cells conditioned media, M2 macrophage monoculture conditioned media, and M2 co-culture with 231 or 468 cells conditioned media. For conditioned media experiments, macrophages were first plated in their respective activation media, before being switched into M0 media and either being co-cultured with cancer cells or remaining in culture alone for 24 hours (as per the conditions listed above). Macrophages and cancer cells were plated at a ratio of 2:1 in 6-well dishes with 1 mL of media. Conditioned media was collected from culture conditions indicated above and added to “naïve” cancer cells (previously untreated cancer cells) for 24 hours prior to the experiment. A conditioned media cancer cell monoculture control was included to account for potential nutrient depletion in conditioned media treatments. This control consisted of naïve cancer cells treated with cancer cell conditioned media (cancer cell monoculture media collected from 24 hour culture prior to experiment).

### Annexin V cell death assay

Fluorescently labelled MDA-MB-231 cells (mito-mEmerald, Addgene, 185596) were cultured with M1 or M2 activated macrophages in direct co-culture in macrophage media + GM-CSF (20 ng/mL) without additional polarization cytokines. After co-culture for 24, 48 or 72 hours, cells were enzymatically dissociated using trypsin-EDTA (25200056, ThermoFisher) and resuspended in serum containing media. Full dead controls used in flow cytometry analysis were prepared by lifting cells by scraping, followed by heating at 55°C for 20 minutes. Dead cells were cooled on ice after incubation. Alive/dead controls of half alive macrophages/half dead macrophages and half alive cancer cells/half cancer cells were included for fluorescence minus one (FMO) - Annexin V controls to determine gates for Annexin V-positivity. Cell samples were centrifuged at 600G for 5 minutes, and pellets were resuspended in BV711-CD11B antibody (macrophage marker; BioLegend, 301344) suspended in flow buffer (1:20 dilution) for 30 minutes on ice in the dark. Following centrifugation, cells were resuspended in Annexin V antibody (A35122, ThermoFisher) diluted in 1X Annexin V binding buffer (556454, BD Biosciences). Annexin V binding buffer was used to dilute the antibody prior to centrifugation and resuspension in flow cytometry buffer. Flow cytometry was performed on the BD LSR Fortessa (5 Lasers: UV, 405, 488, 561, 640). Cancer cells were distinguished from macrophages based on mEmerald-expression. Macrophages were distinguished by BV711-CD11B (1:20)-positive staining. No color controls were included to establish cancer cell+ gating, which was applied to all samples. Gating strategies are outlined in Supplemental Figure 3.

### Macrophage polarization status by flow cytometry

M1 and M2 activated macrophages were plated in monoculture or co-culture with MDA-MB-231 cells in macrophage media + GM-CSF (20 ng/mL) without additional polarization cytokines. Cells were enzymatically lifted with Trypl-E Express Enzyme (1x) (12605028, ThermoFisher) so as not to cleave surface antigen markers. Once lifted, cells were suspended in BV711-CD11B antibody (macrophage marker; BioLegend, 301344, 1:20 dilution) for 30 minutes, to distinguish macrophages from cancer cells in co-culture samples. A macrophage marker for M1 macrophages, anti-human CD80, conjugated to PE-Cyanine7 (305218, BioLegend, 1:20 dilution) and a marker for M2 macrophages, anti-human CD206, conjugated to AlexaFluor 488 (321113, BioLegend, 1:20 dilution) were used to assess polarization status of macrophages at both 24, 48 and 72 hour time points. Macrophages were distinguished from cancer cells based on CD11B positivity, determined by a fluorescent minus one (FMO) CD11B control. FMO controls for macrophage polarization markers were included in these experiments in addition to single color controls for appropriate compensation and gating. CD206+ macrophages were determined based on FMO-CD206 stained M2 activated samples. CD80+ macrophages were similarly determined based on FMO-CD80 stained M1 activated samples. Gating strategy is outlined in Supplemental Figure 2, A-C.

### Statistical analysis

With a minimum expected difference of 30% between conditions, and a standard deviation of 3%, we required at minimum of 3 biological replicates to determine statistical significance at p<0.05. For conditioned media experiments due to a larger standard deviation, we required a minimum of 4 biological replicates. Statistical analyses were done using GraphPad Prism and presented as mean values +/- SEM. Experimental outliers greater than 2 standard deviations from the mean were removed from analysis. Specific tests (2-tailed) and biological replicates are indicated in each figure legend. Biological replicates are indicated as shades of gray on each graph. For all data we assumed gaussian distributions. For flow cytometry data, FlowJo software (version 10.10.0) was used for analysis. For Incucyte assays, the Incucyte Sartorius cell analysis software (version 2021C) was used to measure cell confluence. Image analysis for cell spreading was done using FIJI (version 2.14.0/1.54f).

## Notes

### Competing Interest Statement

The authors have declared no competing interest.

