## Supplemental Figures & Legends for "Macrophage subtypes inhibit breast cancer proliferation in culture"

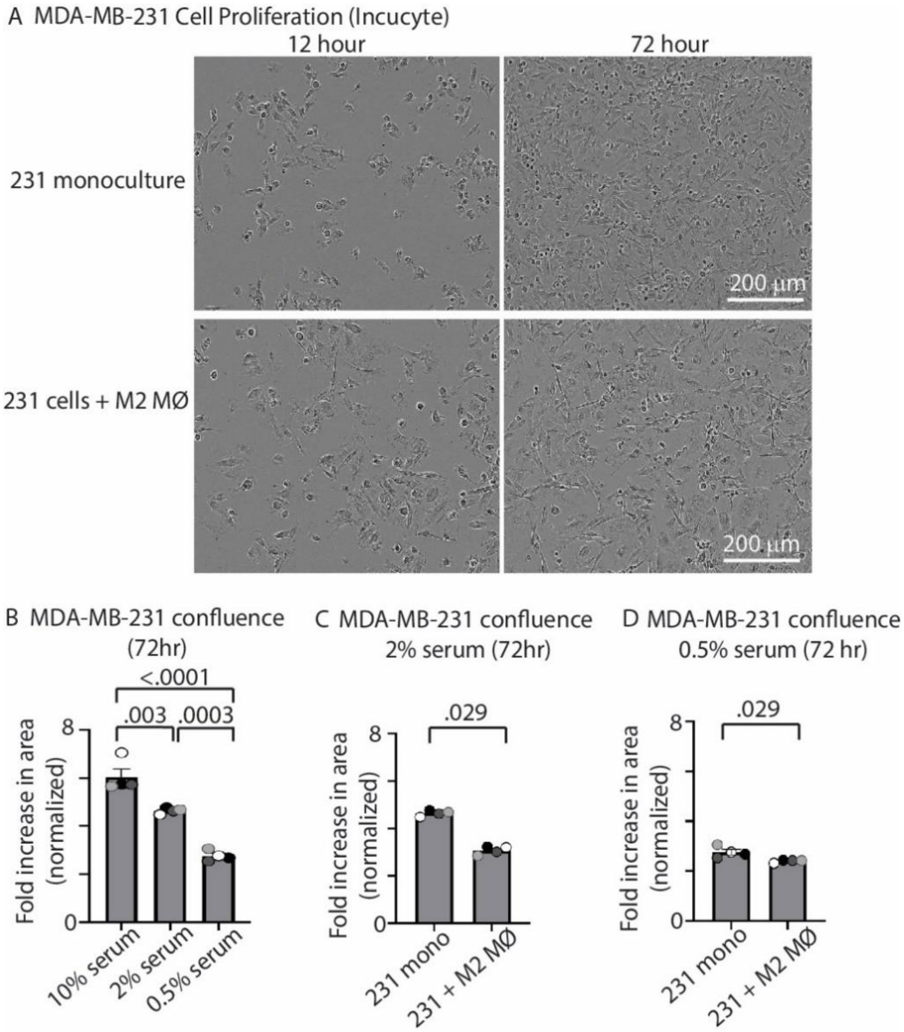


**Supplemental Figure 1. M2-like macrophages inhibit MDA-MB-231 proliferation compared to MDA-MB-231 cells alone.**
(A) Representative stills from a movie of MDA-MB-231 cell proliferation when in monoculture and with M2-like macrophages at 12 and 72 hours in 10% serum conditions. (B) MDA-MB-231 confluence at 72 hours when MDA-MB-231 cells are in 10%, 2%, and 0.5% serum (n=4 independent donors, Tukey's 1-way ANOVA). (C) MDA-MB-231 confluence when MDA-MB-231 cells were plated in monoculture ("231 mono") and when plated with M2-like macrophages ("231 + M2 MΦ") in 2% serum conditions for 72 hours (n=4 independent donors; Mann-Whitney test). (D) MDA-MB-231 confluence when MDA-MB-231 cells were plated in monoculture ("231 mono") and when plated with M2-like macrophages in direct co-culture ("231 + M2 MΦ") in 0.5% serum conditions for 72 hours (n=4 independent donors; Mann-Whitney test).

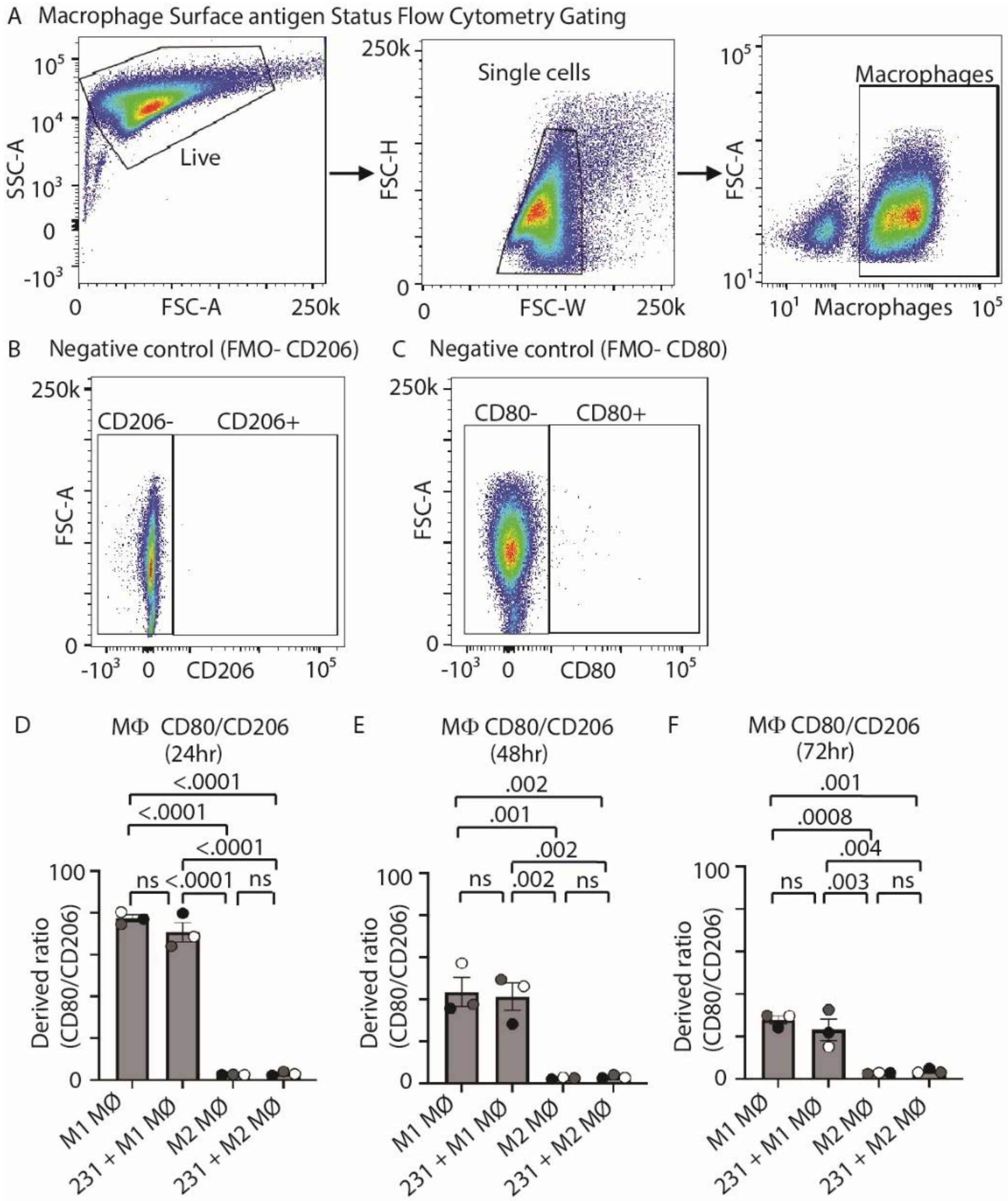


**Supplemental Figure 2. Macrophages retain their differentiation status over time.**
(A) Representative gating for macrophage surface antigen expression. (B) Representative gating for how CD206-positive populations were defined by setting a negative population with FMO-CD206 (see methods) (C). Representative gating for how CD80-positive populations were defined by setting a negative population with FMO-CD80 (see methods). (D-F) CD80/CD206 median derived ratio in M1-like and M2-like macrophages in the following culture conditions: M1-like macrophage monoculture ("M1 MΦ"), MDA-MB-231/M1-like macrophage co-culture ("231 + M1 MΦ"), M2-like macrophage monoculture ("M2 MΦ") and MDA-MB-231/M2-like macrophage co-culture ("231 + M2 MΦ") measured at 24 (D), 48 (E) and 72 (F) hours (n=4 independent donors; Tukey's 1-way ANOVA).

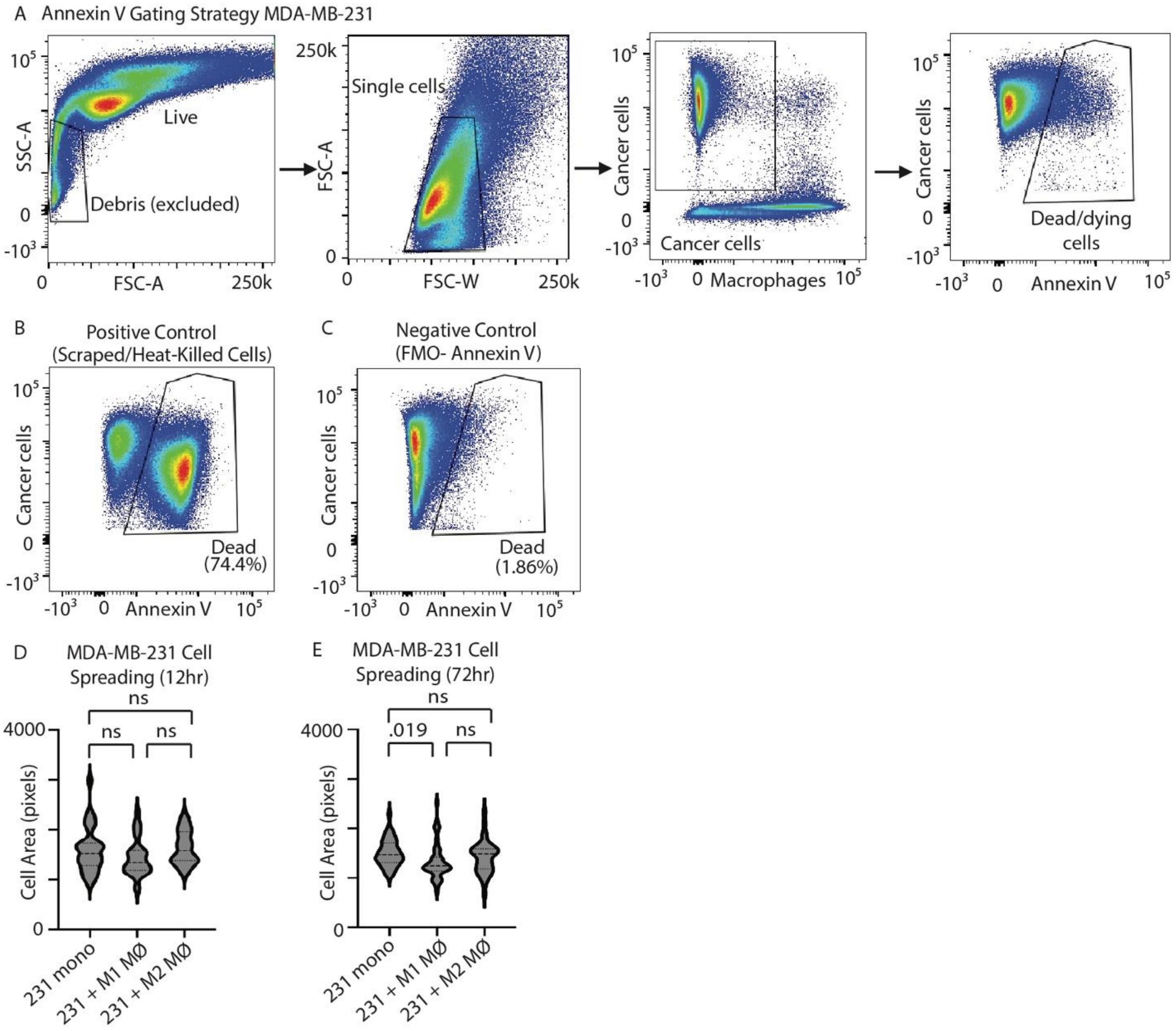


**Supplemental Figure 3. M2-like macrophages do not induce cancer cell death or cell spreading.**

(A) Representative gating strategy for Annexin V quantification. (B) Representative gating for Annexin V+ positive control (heat-killed cells). (C) Representative gating for Annexin V negative control (FMO-Annexin V). (D-E) MDA-MB-231 cell spreading analysis. Cell area is quantified at 12 hours (D) and 72 hours (E) in MDA-MB-231 monoculture ("231 mono"), MDA-MB-231/M1-like macrophage co-culture ("231 + M1 MΦ") and MDA-MB-231/M2-like macrophage co-culture ("231 + M2 MΦ") (12 hours: n=29 cells (231 mono), n=33 cells (231 + M1 mac), n=27 cells (231 + M2 mac); one way-ANOVA). (72 hours: n=44 cells (231 mono), n=56 cells (231 + M1 mac), n=39 cells (231 + M2 mac); Tukey's 1-way ANOVA).

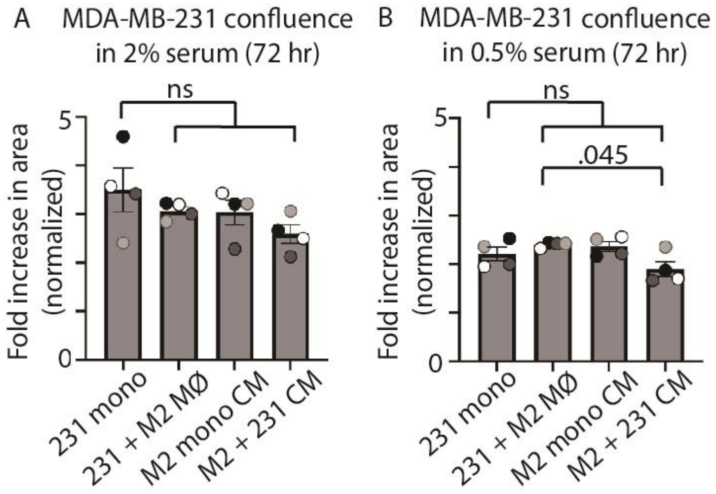


**Supplemental Figure 4. Effects of low serum conditioned media on MDA-MB-231 proliferation.**
(A) MDA-MB-231 cell confluence at 72 hour time point of Incucyte assay in 2% serum when MDA-MB-231 cells are plated in monoculture ("231 mono"), when MDA-MB-231/M2-like macrophages are plated in direct co-culture ("231 + M2 MΦ"), when naïve MDA-MB-231 cells are treated with M2-like macrophage conditioned media ("M2 mono CM") and when naïve MDA-MB-231 cells are treated with MDA-MB-231/M2-like macrophage co-culture CM ("M2 + 231 CM") (B) MDA-MB-231 cell confluence at 72 hour time point of Incucyte assay in 0.5% serum in conditioned media conditions as described in (A). (n=4 independent donors; Tukey's 1-way ANOVA).

**Supplemental Video 1:** MDA-MB-231 cell confluence in monoculture (left) versus M2-like macrophage co-culture (right). Images were taken on the Incucyte at 20x every 4 hours for 72 hours. Scalebar is 200 μm. 2fps.
